# Electrophysiological maturation and increased excitability of human iPSC-derived neurons in *HTR2A* variant-related sleep bruxism

**DOI:** 10.1101/2021.01.26.428254

**Authors:** Avijite Kumer Sarkar, Shiro Nakamura, Kento Nakai, Takahiro Shiga, Yuka Abe, Yurie Hoashi, Tomio Inoue, Wado Akamatsu, Kazuyoshi Baba

**Affiliations:** Department of Prosthodontics, Showa University School of Dentistry, 2-1-1 Kitasenzoku, Ota-ku, Tokyo 145-8515, Japan; Department of Oral Physiology, Showa University School of Dentistry, 1-5-8 Hatanodai, Shinagawa-ku, Tokyo 142-8555, Japan; Center for Genomic and Regenerative Medicine, Juntendo University School of Medicine, 2-1-1 Hongo, Bunkyo-ku, Tokyo 113-8421, Japan

**Keywords:** Sleep bruxism, Human induced pluripotent stem cells, Patch-clamp technique, Intrinsic membrane properties, iPSC-derived neuron maturation, altered excitability

## Abstract

Sleep bruxism (SB) is a sleep-related movement disorder characterized by grinding and clenching of the teeth during sleep. We previously found a significant association between SB and a single nucleotide polymorphism (SNP), rs6313, in the neuronal serotonin 2A receptor gene (*HTR2A*), and established human induced pluripotent stem cell (hiPSC)-derived neurons from SB patients with a genetic variant. To elucidate the electrophysiological characteristics of SB iPSC-derived neural cells bearing a SB-related genetic variant, we generated ventral hindbrain neurons from two SB patients and two unaffected controls and explored the intrinsic membrane properties of these neurons by patch-clamp technique. We found that the electrophysiological properties of the iPSC-derived neurons from the control line mature in a time-dependent manner in long-term cultures. In the early stage of neurogenesis, neurons from two SB lines tended to display shorter action potential (AP) half durations, which led to an increased cell capability of evoked firing. This is the first *in vitro* modelling of SB using disease-specific hiPSCs. The revealed electrophysiological characteristics may serve as a benchmark for further investigation of pathogenic mechanisms underlying SB.

**Summary Statement:** Sleep bruxism patient-specific iPSC-derived neurons with the *HTR2A* variant show altered electrophysiological characteristics, providing the foremost narration of sleep bruxism neurological phenotypes *in vitro* from any species.

## Introduction

Sleep bruxism (SB) is defined as a sleep-related movement disorder characterized by grinding and clenching of the teeth during sleep (Sateia, 2014; Lobbezoo et al., 2013). SB is responsible for a variety of pain and dysfunctional conditions in the orofacial region, such as temporomandibular disorders, abnormal tooth attrition, fracture of teeth or roots, and aggravation of periodontal diseases, which seriously compromise a patient’s quality of life (Beddis et al., 2018).

It is well documented that SB-related masticatory muscle activities occur as part of the partial arousal phenomena, which is associated with transient elevations of central and sympathetic nervous system activity during sleep (Kato et al., 2003; Lavigne et al., 2008), suggesting the possible involvement of central neurotransmitters in the pathogenesis of SB. Several studies reported an association between SB and centrally acting drugs, such as selective serotonin reuptake inhibitor (Ellison and Stanzianin, 1993; Gerber and Lynd, 1998), serotonin 1A receptor agonist (Bostwick and Jaffee, 1999), or alpha-2 adrenergic receptor agonist (Carra et al., 2010) in both positive and negative manners.

Besides, we previously found a significant association of neuronal serotonin 2A receptor gene (*HTR2A*) single nucleotide polymorphism (SNP) rs6313 with SB (Abe et al., 2012). Serotonin is a neurotransmitter involved in many physiological functions in the brain (Guiard and Giovanni, 2015; Lee and Goto, 2018). The serotonin 2A receptor (5-HT2AR) is a serotonin receptor subtype encoded by *HTR2A*. Mutations in this gene have been associated with susceptibility to a variety of neurological diseases. Indeed, rs6313 has been associated with schizophrenia (Inayama et al., 1996; Williams et al., 1997), psychotic symptoms in Alzheimer’s disease (Holmes et al., 1998), certain features of depression (Arias et al., 2001), and sleep breathing disorders (Xu et al., 2014).

Understanding the functional consequences of genetic variants is a critical first step toward appreciating their disease development roles; however, effects of rs6313 on disease pathogenesis have not been elucidated in detail, mainly because of the limited accessibility of the brain. *In vitro* modeling of human diseases using disease-specific hiPSCs can potentially yield dramatic progress in elucidating neurological disease pathogenic mechanisms (Andoh-Noda et al., 2015; Higurashi et al., 2013; Imaizumi et al., 2012).

To construct an *in vitro* disease model using hiPSCs for SB, we previously established a protocol to efficiently induce *HTR2A*-expressing neurons from hiPSCs (Hoashi et al., 2017). Here, we investigated electrophysiological properties of hiPSC-derived neurons, established from two SB patients with the risk allele (C) of *HTR2A* SNP rs6313 (C/C genotype) and two control individuals without the risk allele (T/T genotype), following our previously established protocol (Hoashi et al., 2017). The hiPSCs-derived neurons in the control line exhibited an expected time-dependent maturation of intrinsic membrane properties, as determined through electrophysiological analysis. In the early stage of neurogenesis, the neurons derived from two SB cell lines tended to exhibit altered AP half duration and firing frequency compared to controls. Additionally, the firing frequency and the injected current gain (slope) was 2-fold higher in SB neurons. These results suggest that the excitability of SB hiPSC-derived ventral hindbrain neurons is increased in the early stage of neurogenesis.

To the best of our knowledge, this is the first report of an *in vitro*, cell-level electrophysiological characterization of SB-specific neurons, which may serve as a benchmark for further investigations of pathogenic mechanisms underlying SB. Our newly established SB-specific hiPSCs disease model is expected to be a promising method to elucidate the SB pathological mechanism associated with the genetic variations of 5-HT2AR.

## Results

### hiPSC differentiated neurons exhibit neuronal characteristics

The hiPSC lines were cultured in a Dulbecco’s modified Eagle’s medium (DMEM) to induce a chemically provoked transitional embryoid-body-like state (Fujimori et al., 2017). Since 5-HT2AR neurons are expressed in the raphe nucleus located on the ventral hindbrain (Guiard and Giovanni, 2015); therefore, we generated neurons with the characteristics of the ventral hindbrain region from hiPSCs using previously portrayed differentiation paradigm (Hoashi et al., 2017) with slight modifications (Fig. 1A). To investigate the regional identity of hiPSC-derived neurons, the expression of *NKX2.2* (a ventral brain marker), *EN1* (a midbrain and anterior hindbrain marker), and *HOXC4* (a posterior hindbrain and a spinal cord marker) were analyzed in hiPSCs (C2 line) at days *in vitro* (DIV) 7 (Fig. 1B). Samples treated with 1 μM retinoic acid (RA), in the absence of sonic hedgehog (Shh) and purmorphamine (PM) treatment, were used as controls. Although cells expressed significantly higher levels of *NKX2.2* and *EN1*, compared to the control, they expressed significantly lower levels of *HOXC4*, suggesting that they had successfully acquired ventral hindbrain identity.

**Figure 1.**
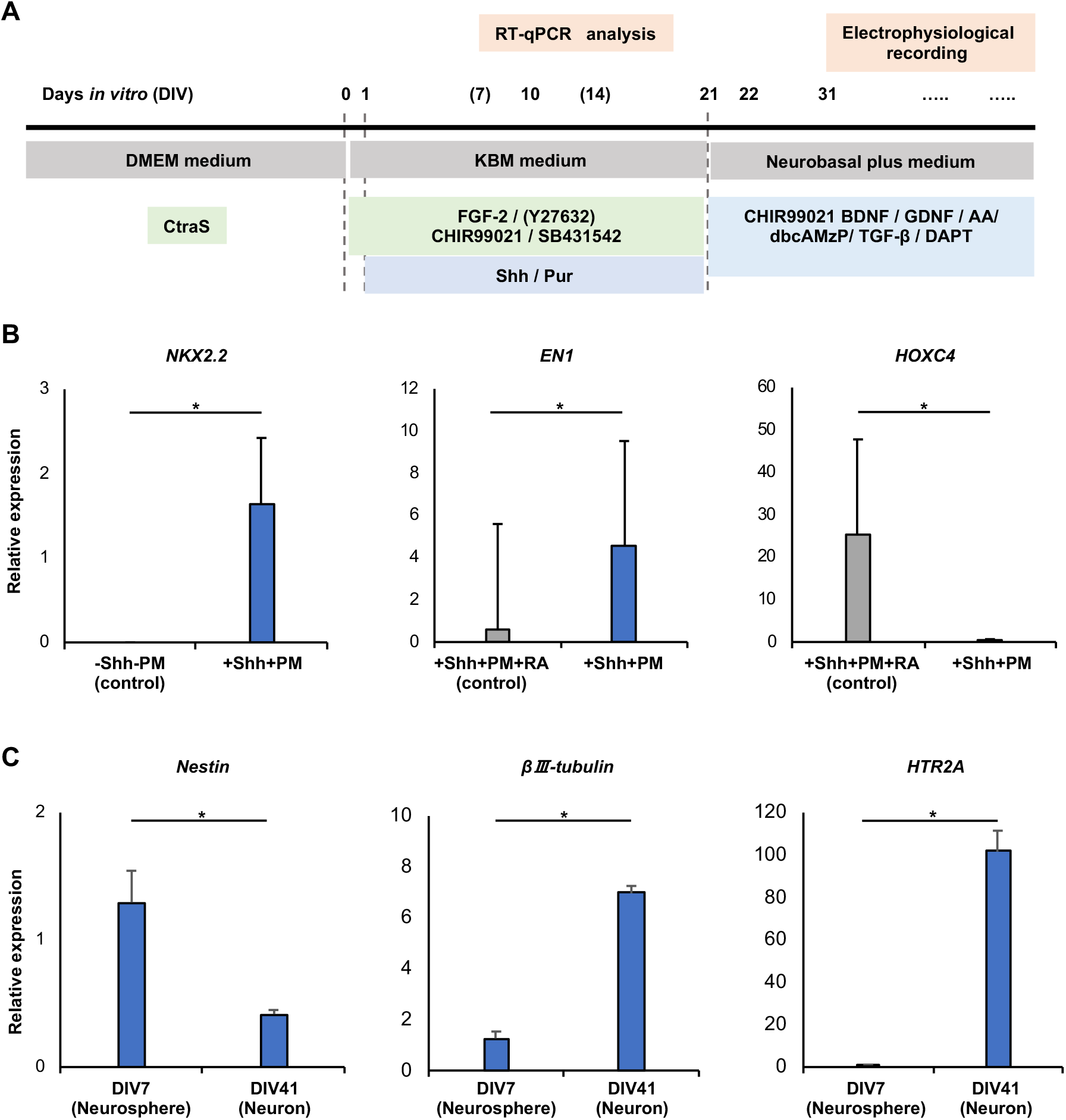
Differentiated neurons from hiPSCs exhibited neuronal characteristics. (A) Schematic overview of the differentiation protocol, including time points for RT-qPCR and electrophysiological analyses. (B) Statistical summary of *NKX2.2* (a ventral brain marker), *EN1* (a midbrain and anterior hindbrain marker), and *HOXC4* (a posterior hindbrain and a spinal cord marker) gene expression in hiPSCs at DIV7. Number of samples assessed: +Shh+PM; *n*=5 and control; *n*=4. (C) Gene expression summary of the neural progenitor cell marker, *Nestin*, and neuronal markers, *βIII-tubulin*, and *HTR2A*, in hiPSCs at DIV7 and 41 (*n*=5). Data are expressed as mean ± s.e.m. \**p*<0.05 (nonparametric Kruskal–Wallis test).

To follow the generation of hiPSC-derived neurons, RT-qPCR analyses were performed on DIV7 and 41 of differentiation. At DIV7, cells expressed significantly higher levels of *Nestin*, suggesting the successful neural patterning of hiPSCs (Fig. 1C). In contrast, *βIII-tubulin* and *HTR2A* expression levels considerably increased at DIV41, indicating the successful neuronal differentiation of hiPSCs expressing 5-HT2AR (Fig. 1C).

### Time-dependent maturation of the intrinsic membrane properties of hiPSC-derived ventral hindbrain neurons in long-term cultures

Whole-cell patch-clamp recordings were performed on control (C2) iPSC-derived ventral hindbrain neurons at DIV 31–51, 52–71, 72–91, and 92–111 in current-clamp mode to trace the functional maturity of their intrinsic membrane properties over the course of neurogenesis for the expected electrophysiological maturation of differentiated neurons (Fig. 2A).

**Figure 2.**
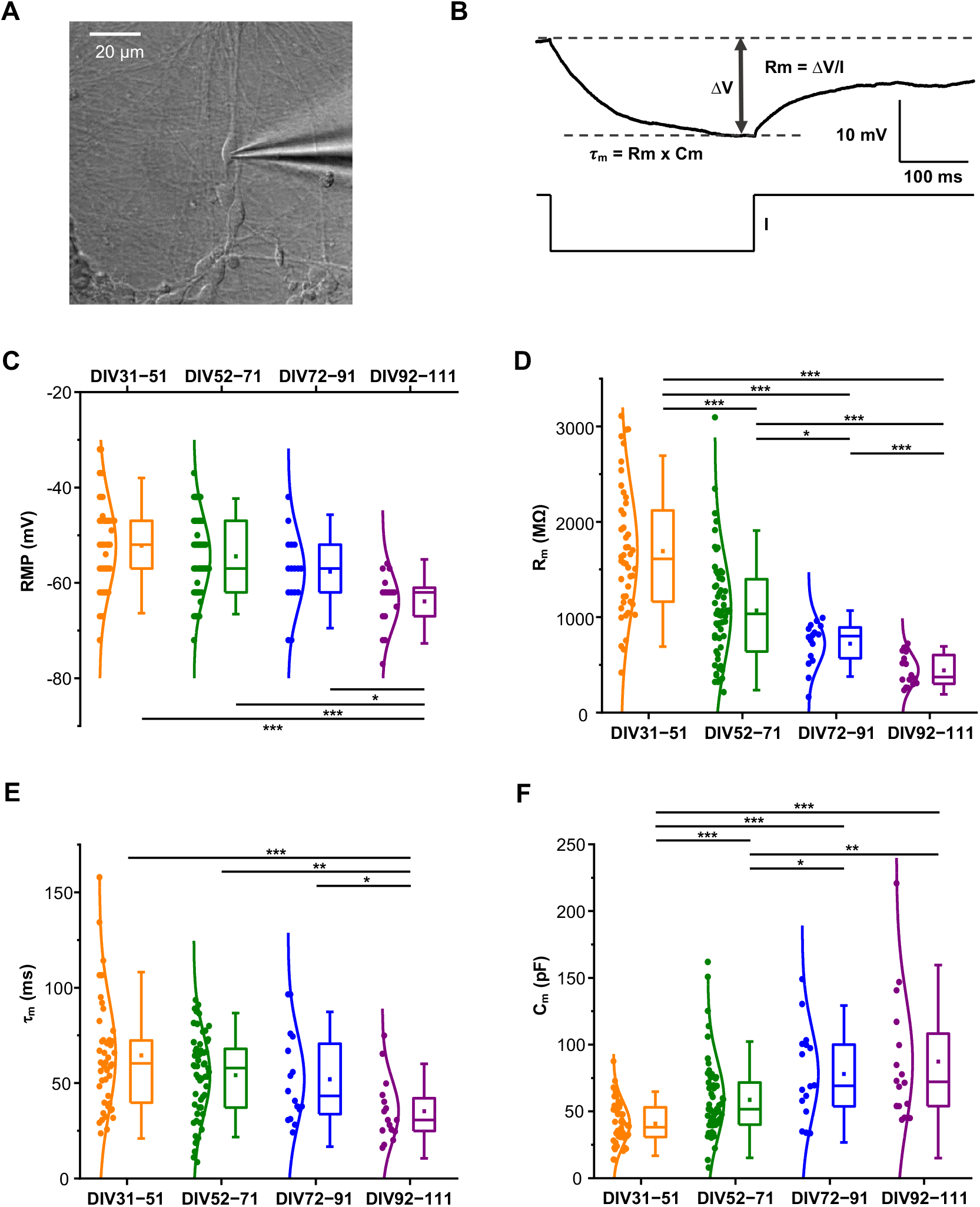
hiPSC-derived ventral hindbrain neurons changed passive membrane properties in a time-dependent manner in the unaffected control (C2) cell line. (A) DIC image depicting a typical hiPSC-derived neuron being approached by a patching pipette during electrophysiological recordings at 31–51 days *in vitro* (DIV) of neurogenesis. (B) Parameters extracted to interpret the passive membrane properties of hiPSC-derived neurons. The membrane resistance (R_m_), the time constant (τ_m_) and the membrane capacitance (C_m_) were calculated based on the voltage response (top trace) of the cell to a small step of hyperpolarizing current (bottom trace). (C–F) Boxplot and whisker plots containing raw data and distribution curves show a statistical overview of (C) the resting membrane potential (RMP), (D) the membrane resistance (R_m_), (E) the time constant (τ_m_), and (F) the membrane capacitance (C_m_) of differentiated neurons at the time points of DIV31–51, DIV52–71, DIV72–91, and DIV92–111 time points of neurogenesis. Color coding: DIV31–51, orange; DIV52–71, olive; DIV72–91, blue; DIV92–111, purple. Sample size: *n*=46 at DIV31–51, *n*=59 at DIV52–71, *n*=16 at DIV72–91, and *n*=16 at DIV92–111. Boxes represent the 25–75 percentile of data distribution. Squares inside the boxes and whiskers indicate mean values and standard deviations, respectively. \**p*<0.05, \*\**p*<0.01, and \*\*\**p*<0.001 (nonparametric Mann–Whitney U test for pairwise comparisons).

To determine both electrophysiological maturity and heterogeneity between different time points, we started by quantifying and comparing the passive membrane properties of differentiated cells. Passive properties, including membrane resistance (R_m_), time constant (τ_m_), and membrane capacitance (C_m_), were obtained from voltage responses to small hyperpolarizing current pulses applied around the resting membrane potential (RMP) (Fig. 2B). Firstly, over the course of neurogenesis, the RMP became dynamically more hyperpolarized (RMP, DIV31–51 −52.2 ± 1.4 mV, *n*=46; DIV52–71 −54.4 ± 1.0 mV, *n*=59; DIV72–91 −57.6 ± 2.0 mV, *n*=16; DIV92–111 −63.9 ± 1.5 mV, *n*=16; *p*=0.0001; Kruskal–Wallis test; Fig. 2C). Secondly, there was a 74% decrease in the R_m_ from DIV31–51 to DIV92–111, highlighting the progressive addition of ion channels to the plasma membrane (R_m_, DIV31–51 1692.2 ± 98.4 MΩ, *n*=46; DIV52–71 1071.1 ± 72.6 MΩ, *n*=59; DIV72–91 722.2 ± 57.6 MΩ, *n*=16; DIV92–111 442.6 ± 41.9 MΩ, *n*=16; *p*<0.0001; Kruskal–Wallis test; Fig. 2D). Another passive property of the membrane, the τ_m,_ also markedly reduced by DIV92-111 (τ_m_, DIV31–51 64.5 ± 4.3 ms, *n*=46; DIV52–71 54.2 ± 2.8 ms, *n*=59; DIV72–91 52.0 ± 5.9 ms, *n*=16; DIV92–111 35.3 ± 4.1 ms, *n*=16; *p*=0.001; Kruskal–Wallis test; Fig. 2E). Thirdly, the C_m_ sharply increased, – almost 2-fold – between DIV31–51 and DIV92–111 (C_m_, DIV31–51 40.8 ± 2.4 pF, *n*=46; DIV52–71 58.7 ± 3.8 pF, *n*=59; DIV72–91 78.1 ± 8.5 pF, *n*=16; DIV92–111 87.4 ± 12.0 pF, *n*=16; *p*<0.0001; Kruskal–Wallis test; Fig. 2F).

The more negative RMP, lower Rm, shorter τ_m_, and larger C_m_ over DIV111 reflected the general trends documented during neuronal maturation (Kang et al., 2017; Kopach et al., 2018; Lam et al., 2017; Prè et al., 2014; Weick, 2016). Collectively, these four parameters were determinant of hiPSC-derived neuronal maturation in long-term cultures.

Next, we assessed the active membrane properties of generated neurons. Differentiated neurons could generate spontaneous and evoked action potentials (APs) during extended periods of neurogenesis (Fig. 3A,C). The proportion of neurons firing spontaneous APs tended to increase during maturation (DIV31–51 54.2%, 13/24; DIV52–71 60%, 24/40; DIV72–91 63.6%, 7/11; DIV92–111 85.7%, 6/7; *p*=0.460; Chi-squared test; Fig. 3B). Consistent with the increased proportion of spontaneous AP firing, neurons evolved their capability to high-frequency discharge in response to an injection of a series of depolarizing current pulses (Fig. 3C). To further assess neuronal maturity, we utilized a ranking framework classifying neurons into three types depending on their overshooting amplitude of APs and firing frequency (Fig. 3D; see Materials and Methods). This analysis revealed that the proportion of Type 3 neurons which fired APs above 0 mV at a maximal frequency of 10 Hz or more, sharply increased over time (DIV31–51 63.0%, 29/46; DIV52–71 83.0%, 49/59; DIV72–91 93.8%, 15/16; DIV92–111 100%, 16/16; *p*=0.003; Chi-squared test; Fig. 3E). To illustrate the developmental evolution of AP shape, we measured four properties from the first evoked AP in response to minimum current injection: AP rheobase (R_h_), AP threshold, AP amplitude, and AP half duration (Fig. 4A). Both the R_h_ (R_h_, DIV31–51 19.3 ± 1.9 pA, *n*=46; DIV52–71 24.9 ± 2.1 pA, *n*=59; DIV72–91 28.7 ± 3.5 pA, *n*=16; DIV92–111 46.9 ± 7.5 pA, *n*=16; *p*<0.0001; Kruskal– Wallis test; Fig. 4B) and amplitude (amplitude, DIV31–51 80.9 ± 2.6 mV, *n*=46; DIV52–71 89.0 ± 2.0 mV, *n*=59; DIV72–91 93.5 ± 3.9 mV, *n*=16; DIV92–111 110.0 ± 2.3 mV, *n*=16; *p*<0.0001; Kruskal–Wallis test; Fig. 4C) remarkably increased during neuronal development. Thirdly, the AP threshold (threshold, DIV31–51 −46.0 ± 0.8 mV, *n*=46; DIV52–71 −47.9 ± 0.5 mV, *n*=59; DIV72–91 −49.6 ± 1.4 mV, *n*=16; DIV92–111 −53.8 ± 1.0 mV, *n*=16; *p*<0.0001; Kruskal–Wallis test; Fig. 4D) became more hyperpolarized over time. Furthermore, AP half duration (half duration, DIV31–51 5.9 ± 0.6 ms, *n*=46; DIV52–71 4.5 ± 0.4 ms, *n*=59; DIV72– 91 2.9 ± 0.2 ms, *n*=16; DIV92–111 1.8 ± 0.1 ms, *n*=16; *p*<0.0001; Kruskal–Wallis test; Fig. 4E) shortened significantly at DIV92–111 after differentiation was induced.

**Figure 3.**
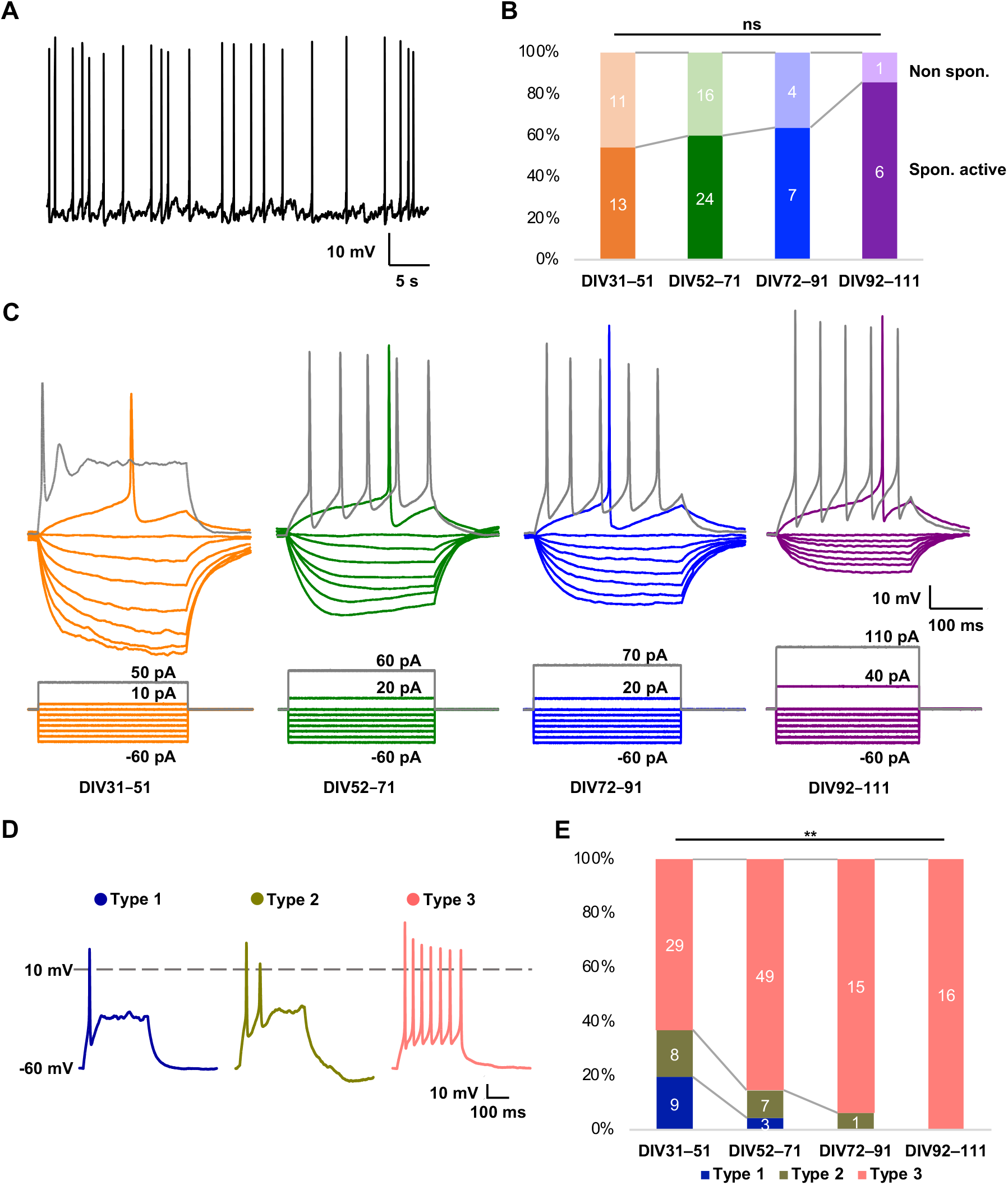
AP firing of human cells in the control (C2) line mature over time. (A) Example traces of spontaneous APs from a spontaneously active hiPSC-derived neuron. (B) Proportion of neurons firing spontaneous APs at DIV31–51, DIV52–71, DIV72–91, and DIV92–111 of neurogenesis. Number of total cells analyzed: *n*=24 at DIV31–51, *n*=40 at DIV52–71, *n*=11 at DIV72–91, and *n*=7 at DIV92–111. Chi-squared test. ns = nonsignificant. (C) Representative examples of the typically evoked firing (upper panel) of neurons at DIV31–51, 52–71, 72–91, and 92–111 in response to 300 ms-long consecutive steps of square current pulses (lower panel). Membrane potential was held at −60 mV. Note the high-frequency discharge evoked by a current of greater stimulating intensity in neurons at DIV52–71, 72–91, and 92–111, compared to the single AP spike elicited at DIV31–51 (gray traces). (B,C) Color coding: DIV31–51, orange; DIV52–71, olive; DIV72–91, blue; DIV92–111, purple. (D) Representative traces of typical heterogeneous neuronal responses to optimal depolarizing current pulses for 300 ms (membrane potential clamped at −60 mV). Depending on the overshooting (0 mV) amplitude of APs and the firing frequency, the heterogeneous states of generated neurons were categorized into a spectrum of three AP types. (E) Diagram representing the proportion of neuronal maturity classes at DIV31–51, DIV52–71, DIV72–91, and DIV92–111 of neurogenesis. Number of total cells assessed: *n*=46 at DIV31–51, *n*=59 at DIV52–71, *n*=16 at DIV72–91, and *n*=16 at DIV92–111. \*\**p*<0.01 (Chi-squared test). (D,E) Color coding: Type 1 neurons, royal; Type 2 neurons, dark yellow; Type 3 neurons, rose.

**Figure 4.**
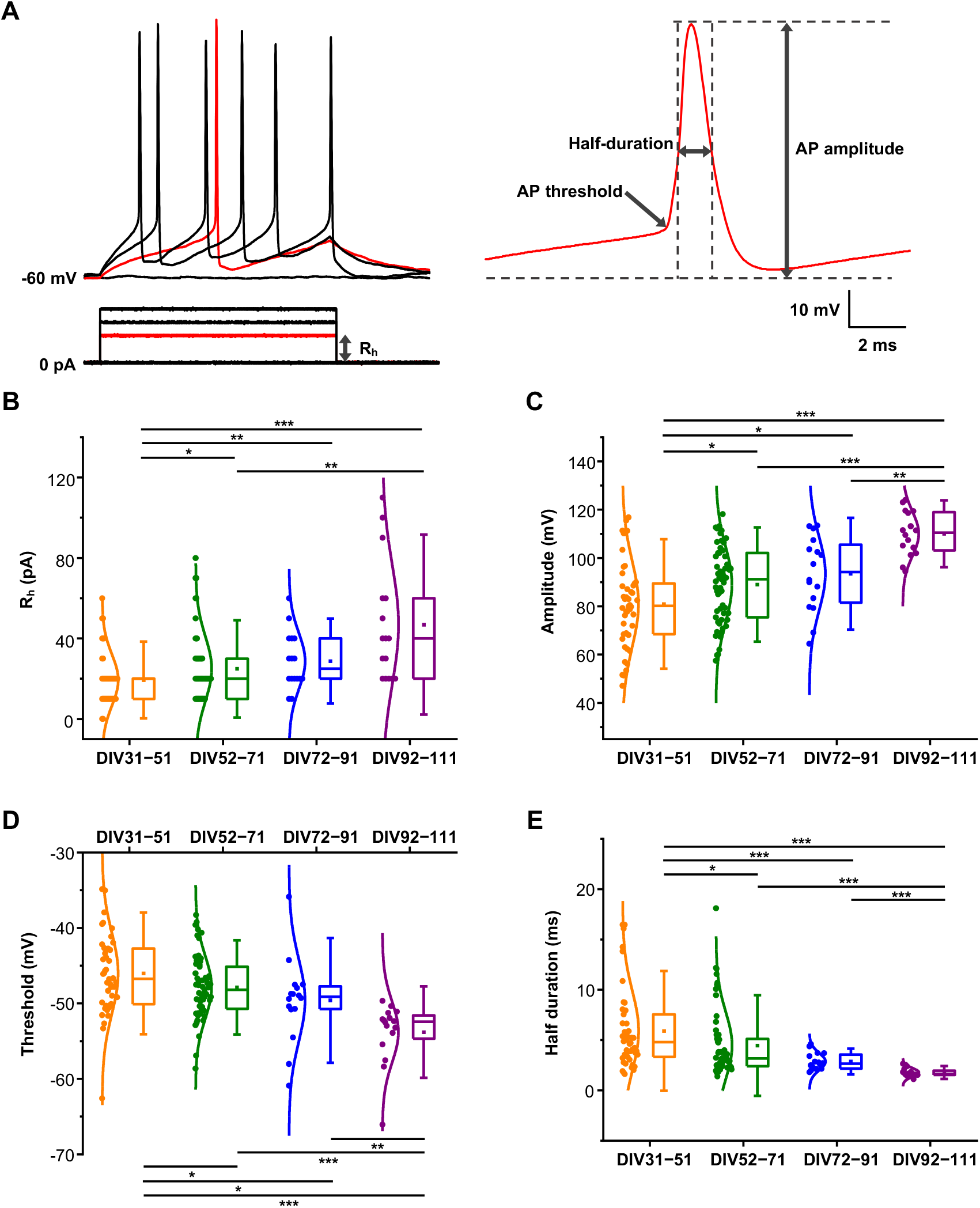
AP characteristics of control (C2) hiPSC-derived neurons evolved during maturation. (A) Example traces of a typical voltage recording in response to a minimum depolarizing current step depicting the parameters calculated to characterize the APs in human cells. Four parameters were calculated, including rheobase (R_h_) (left) and threshold, amplitude, and half duration (right). (B-E) Box and whisker plots showing the statistical summary of (B) the R_h_, (C) the threshold (D) the amplitude, and (E) the half duration in hiPSC-derived neurons at DIV31–51, DIV52–71, DIV72–91, and DIV92–111 of neurogenesis. Color coding: DIV31–51, orange; DIV52–71, olive; DIV72–91, blue; DIV92–111, purple. Sample size: *n*=46 at DIV31–51, *n*=59 at DIV52–71, *n*=16 at DIV72–91, and *n*=16 at DIV92–111. Boxes represent the 25–75 percentile of data distribution. Squares inside the boxes and whiskers indicate mean values and standard deviations, respectively. \**p*<0.05, \*\**p*<0.01, and \*\*\**p*<0.001 (nonparametric Mann–Whitney U test for pairwise comparisons).

Altogether, long-term cultures of hiPSC-derived neurons showed clear signs of functional maturation over the course of DIV111, including a higher proportion of spontaneously active, and type 3 neurons, an increased AP R_h_, a greater AP amplitude, a hyperpolarized AP threshold, and a shorter AP half duration; each of which are key aspects enabling differentiated neurons to be able to form neural communication (Lam et al., 2017).

### SB iPSC-derived neurons show altered intrinsic membrane properties

As the intrinsic membrane properties of control (C2) neurons mature over time, we obtained patch-clamp recordings from iPSC-derived neurons of two SB patients (SB1 and SB2) and a different unaffected control (C1) individual at DIV31–51, to determine whether there were alterations in SB neurons. Firstly, we found no significant difference in the RMP of iPSC-derived neurons from the two SB lines, as compared to the two age-matched control lines (C1, −57.2 ± 3.1 mV, *n*=8; C2, −52.2 ± 1.4 mV, *n*=46; SB1, −56.6 ± 2.8 mV, *n*=12; SB2, −56.0 ± 1.9 mV, *n*=20, *p*=0.188; Kruskal–Wallis test; Fig. 5A). Secondly, one of the other passive membrane properties, Rm, was also not substantially different in neurons between SB and the age-matched controls (C1, 1850.6 ± 277.6 MΩ, *n*=8; C2, 1692.2 ± 98.4 MΩ, *n*=46; SB1, 1280.4 ± 135.5 MΩ, *n*=12; SB2, 1343.7 ± 145.7 MΩ, *n*=20; *p*=0.061; Kruskal–Wallis test; Fig. 5B). Thirdly, there was no noticeable distinction observed in τ_m_ between SB and control neurons (C1, 69.2 ± 8.1 ms, *n*=8; C2, 64.5 ± 6.8 ms, *n*=46; SB1, 60.1 ± 8.2 ms, *n*=12; SB2, 54.2 ± 5.3 ms, *n*=20; *p*=0.374; Kruskal–Wallis test; Fig. 5C). In addition, the C_m_ in SB neurons was not different from those in control neurons (C1, 39.3 ± 2.6 pF, *n*=8; C2, 40.8 ± 2.4 pF, *n*=46; SB1, 47.0 ± 3.4 pF, *n*=12; SB2, 45.8 ± 4.8 pF, *n*=20; *p*=0.377; Kruskal–Wallis test; Fig. 5D).

**Figure 5.**
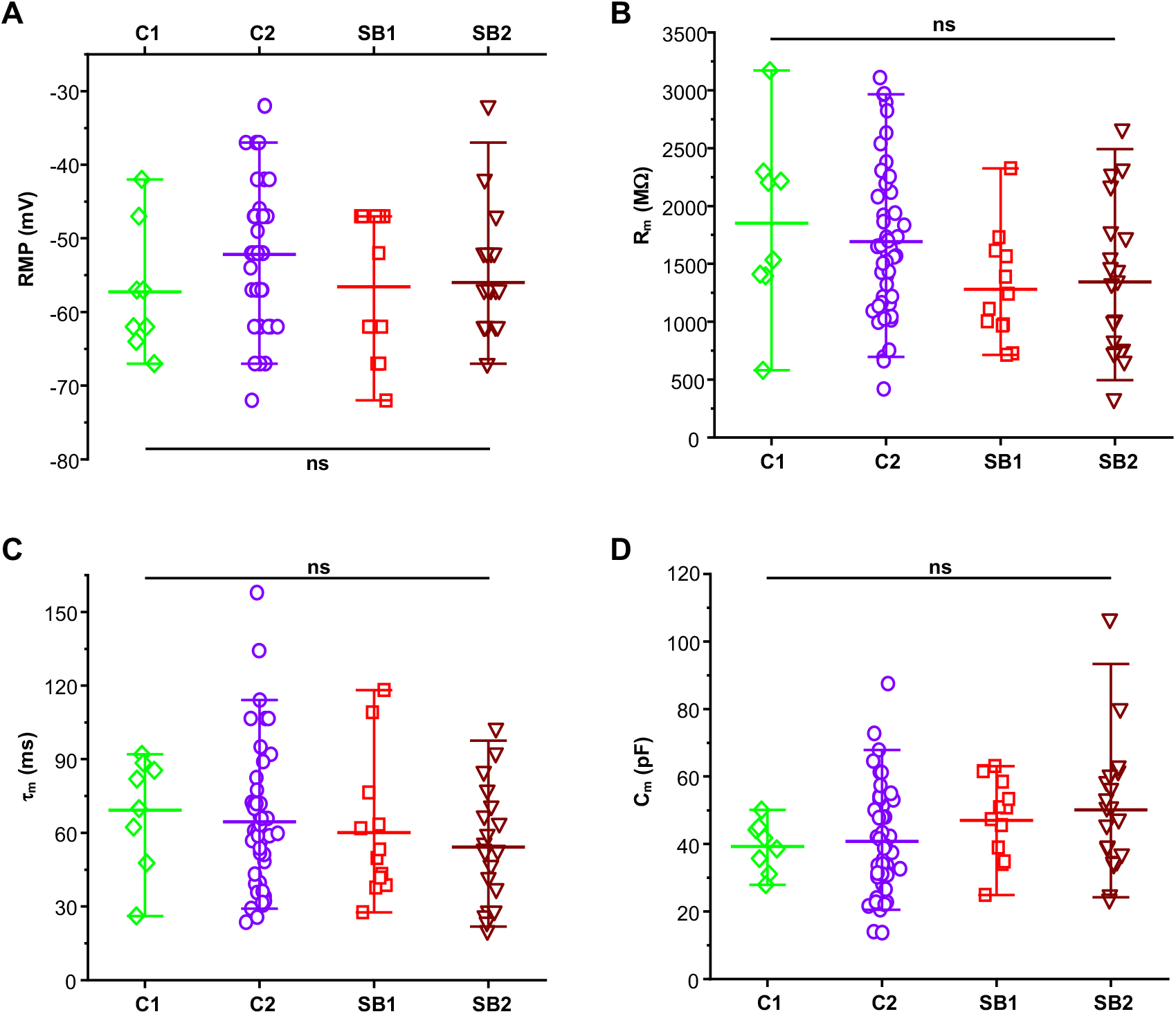
Passive membrane properties of hiPSC-derived neurons in SB cell lines were similar to control cell lines at DIV31–51. (A-D) Scatter plots containing raw data demonstrate the statistical summary of (A) the resting membrane potential (RMP), (B) the membrane resistance (R_m_), (C) the time constant (τ_m_), and (D) the membrane capacitance (C_m_) of hiPSC-derived neurons in unaffected control (C1 and C2) and two SB patient (SB1, SB2) cell lines at DIV31–51. Color and symbol coding: C1, green diamond; C2, violet circle; SB1, red square and SB2, wine down-pointing triangle. Sample size: *n*=8 in C1, *n*=46 in C2, *n*=12 in SB1 and *n*=20 in SB2. Lines and whiskers represent mean values and the 5–95 percentile, respectively. Nonparametric Kruskal–Wallis test, ns = nonsignificant.

Next, we examined AP firing at a holding voltage of −60 mV in response to a 300 ms-long current stimulus. Firstly, SB neurons showed a significantly larger proportion of cells expressing Type 3 firing (C1, 50%, 4/8; C2, 63%, 29/46; SB1, 91.7%, 11/12; SB2, 95%, 19/20; *p*=0.010; Chi-squared test; Fig. 6A). In addition, current steps of the same intensity generated a higher number of spikes in SB, compared to age-matched control neurons (Fig. 6B,C). The increased firing frequency of SB neurons was statistically significant for current steps of 50, 60, 70, and 80 pA (Fig. 6C and Table 1). Next, frequency-current intensity data from C1 and C2, and SB1 and SB2 were pooled as control and SB, respectively, and fitted to a linear regression (Fig. 6D). As shown in Fig. 6D, upon increasing stimulus intensity, neurons from pooled control cell lines reached a firing plateau with stimuli ≥ 60 pA; for this reason, data were fitted to a linear regression up to 60 pA. Both control and SB neurons exhibited good fit (*r*>0.9); the SB cell group showed a significantly higher gain than the control (0.14 ± 0.02, *n*=54 in control, versus 0.30 ± 0.03, *n*=32 in SB; *p*=0.0001; Kruskal–Wallis test; Fig. 6E,F). The mean gain in SB neurons was 2-fold higher than in control neurons. Secondly, the AP rheobase did not differ between controls and SB neurons (C1, 13.7 ± 2.6 pA, *n*=8; C2, 19.3 ± 1.9 pA, *n*=46; SB1, 15.0 ± 3.4 pA, *n*=12 and SB2, 18.0 ± 2.1 pA, *n*=20; *p*=0.237; Kruskal– Wallis test; Fig. 7A). In addition, the APs amplitudes in SB neurons did not different from those in control neurons (C1, 82.1 ± 3.6 mV, *n*=8; C2, 80.9 ± 2.6 mV, *n*=46; SB1, 84.7 ± 3.2 mV, *n*=12 and SB2, 86.0 ± 3.8 mV, *n*=20; *p*=0.671; one-way ANOVA test; Fig. 7B). Thresholds, another active membrane property of membranes, were not fundamentally different between neurons from SB and control lines; however, SB2 displayed a significantly depolarized threshold in compared to C1 neurons (C1, −50.7 ± 0.8 mV, *n*=8 vs. SB1, −48.3 ± 1.0 mV, *n*=12; *p*=0.123 vs. SB2, −48.2 ± 0.6 mV, *n*=20, *p*=0.022 and C2, −46.0 ± 0.8 mV, *n*=46 vs. SB1, −48.3 ± 1.0 mV, n=12, *p*=0.120 vs. SB2, −48.2 ± 0.6 mV, *n*=20, *p*=0.070; Mann– Whitney U test; Fig. 7C). Furthermore, AP half durations were significantly shorter in the two SB cell lines, compared with the two control lines (5.9 ± 0.5 ms, *n*=8 in C1 vs. 3.3 ± 0.4 ms, *n*=12 in SB1; *p*=0.003 vs. 3.8 ± 0.4 ms, *n*=20 in SB2; *p*=0.003 and 5.9 ± 0.6 ms, *n*=46 in C2 vs. 3.3 ± 0.4 ms, *n*=12 in SB1; *p*=0.016 vs. 3.8 ± 0.4 ms, *n*=20 in SB2; *p*=0.038; Mann–Whitney U test; Fig. 7D,E).

**Table 1.** hiPSC-derived neurons exhibit AP potential firing at the early stage of neurogenesis. Data are expressed as mean ± s.e.m. (number of the cells recorded). AP firing frequencies were evoked using 300 ms-long square depolarizing current pulses at 10, 20, 30, 40, 50, 60, 70, and 80 pA; APs were measured per second. Statistical analysis was performed using ANOVA with Tukey’s post-hoc test and nonparametric Mann–Whitney U test for pairwise comparisons.

| Injected current | Firing frequency [Hz] |  |  |  | *p*-value |  |  |  |
| --- | --- | --- | --- | --- | --- | --- | --- | --- |
|  | C1 | C2 | SB1 | SB2 | C1 vs. SB1 | C1 vs. SB2 | C2 vs. SB1 | C2 vs. SB2 |
| 10 pA | 6.7 ± 1.8 (8) | 3.6 ± 0.7 (46) | 3.3 ± 0.7 (12) | 4.5 ± 1.3 (20) | 0.129 | 0.331 | 0.371 | 0.602 |
| 20 pA | 9.6 ± 2.0 (8) | 7.3 ± 0.8 (46) | 10.6 ± 1.7 (12) | 9.0 ± 1.8 (20) | 0.781 | 0.816 | 0.135 | 0.580 |
| 30 pA | 10.4 ± 1.3 (8) | 8.6 ± 0.9 (46) | 14.7 ± 2.3 (12) | 13.7 ± 1.9 (20) | 0.135 | 0.518 | 0.010 | 0.030 |
| 40 pA | 11.3 ± 1.1 (8) | 10.3 ± 1.0 (46) | 16.7 ± 2.6 (12) | 16.2 ± 1.9 (20) | 0.101 | 0.256 | 0.022 | 0.009 |
| 50 pA | 10.4 ± 1.2 (8) | 11.5 ± 1.1 (46) | 18.3 ± 2.6 (12) | 17.7 ± 1.8 (20) | 0.046 | 0.016 | 0.017 | 0.004 |
| 60 pA | 10.4 ± 1.3 (8) | 11.7 ± 1.1 (46) | 20.6 ± 3.0 (12) | 18.7 ± 1.8 (20) | 0.017 | 0.005 | 0.006 | 0.001 |
| 70 pA | 10.4 ± 1.6 (8) | 11.2 ± 1.0 (46) | 22.5 ± 3.5 (12) | 19.0 ± 2.0 (20) | 0.016 | 0.013 | 0.002 | 0.001 |
| 80 pA | 10.0 ± 1.3 (8) | 10.1 ± 0.9 (46) | 23.9 ± 3.7 (12) | 20.2 ± 2.4 (20) | 0.005 | 0.007 | < 0.001 | < 0.001 |

**Figure 6.**
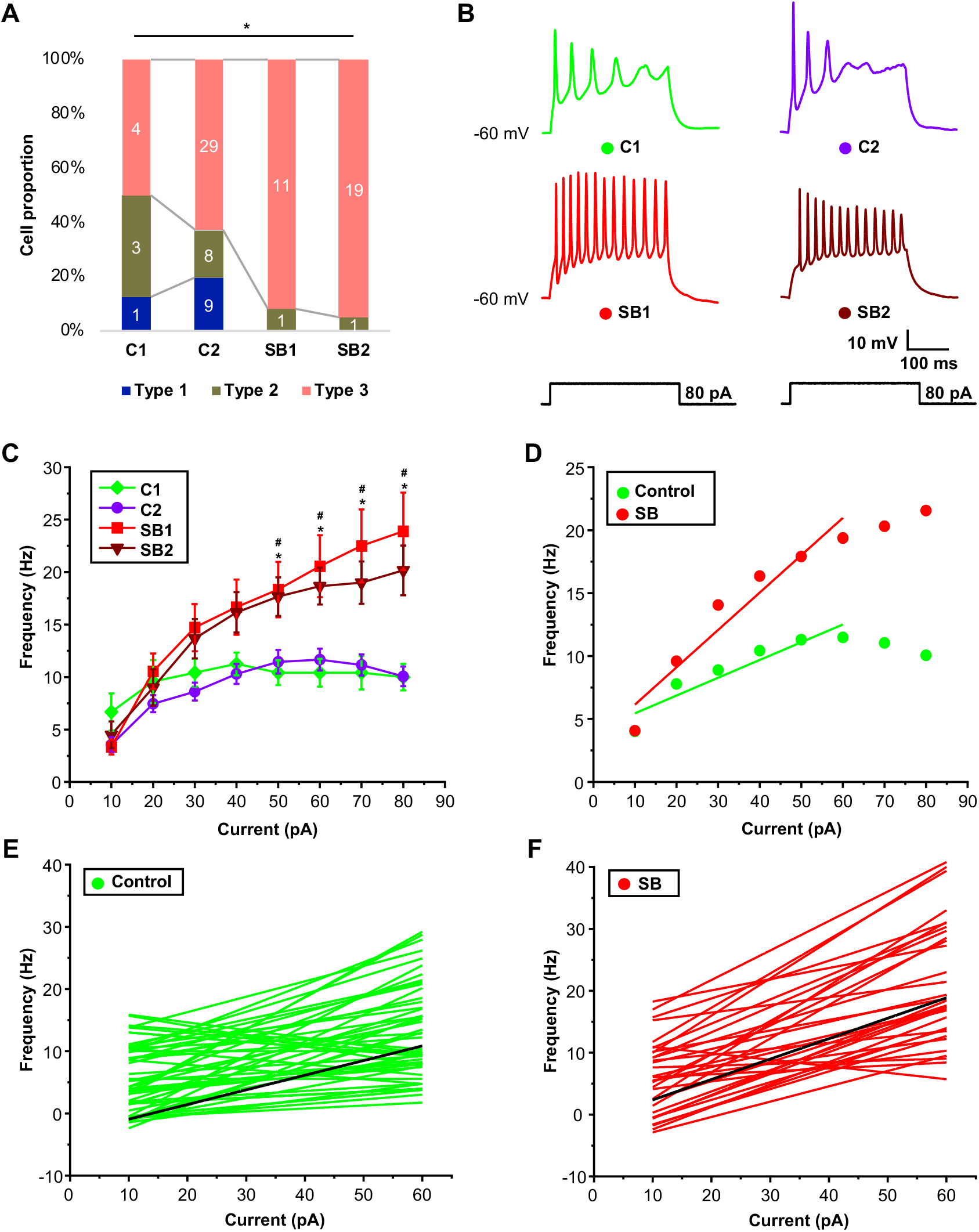
SB iPSC-derived neurons showed altered firing patterns in the early stage of neurogenesis. (A) Proportion of cells belonging to AP type classifications for neurons derived from two control (C1 and C2) and two SB patient (SB1 and SB2) hiPSC lines. Note that nearly all neurons from SB lines exhibited Type 3 APs (overshooting 0 mV at a maximal frequency of 10 Hz or more) at DIV31–51. Color coding: Type 1 neurons, Royal; Type 2 neurons, dark yellow; Type 3 neurons, rose. \**p*<0.05 (Chi-squared test). (B) Examples of APs in response to an 80 pA depolarizing current injection applied for 300 ms (bottom panel) in C1 (left, green), C2 (right, violet) (top panel), SB1 (left, red) and SB2 (right, wine) (middle panel) neurons. (C) Firing frequency-current plot for C1 (*n*=8, green), C2 (*n*=46, violet), SB1 (*n*=12, red), and SB2 (*n*=20, wine) hiPSC-derived neurons. Data are presented as mean ± s.e.m. Statistical analysis was performed using a nonparametric Mann–Whitney U test for pairwise comparisons. * indicates significant differences between C1, SB1, and SB2; # indicates significant differences between C2, SB1, and SB2. Detailed data for each group are presented in Table 1. (D) Plot of the frequency versus injected current relationship for pooled data from control neurons (green) and SB neurons (red). The gain corresponds to a linear regression. Control: Pearson’s *r*=0.94, *p*=0.006, *R*^2^=0.85. SB: Pearson’s *r*=0.96, *p*=0.002, *R*^2^=0.90. (E,F) Linear relationships between injected current and firing frequency for pooled (E) control (green) and (F) SB (red) groups of hiPSC-derived neurons. Black lines correspond to the mean fit for each group.

**Figure 7.**
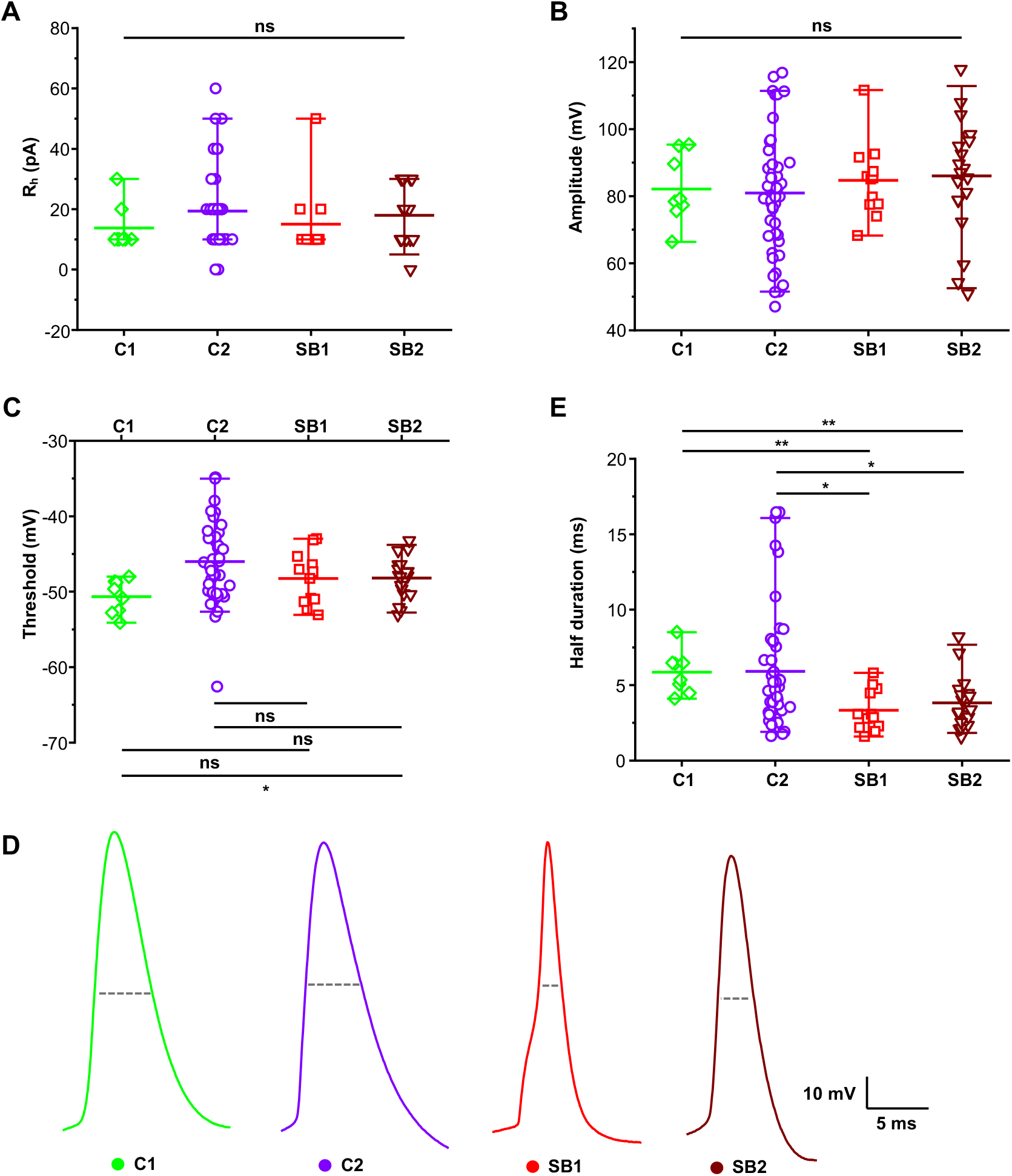
The AP half duration differed between control and SB cell lines. (A,B) Scatter plots containing raw data present a statistical summary of (A) the rheobase (R_h_) and (B) the amplitude of hiPSC-derived neurons in two control (C1 and C2), and two SB (SB1 and SB2) cell lines. Nonparametric Kruskal–Wallis test. ns = nonsignificant. (C) Threshold of hiPSC-derived neurons in two control and two SB lines. \**p*<0.05 (nonparametric Mann–Whitney U test for pairwise comparisons). ns = nonsignificant. (D) Examples of the first individual APs, as evoked in response to minimal current injection, from the hiPSC-derived neurons in C1 (green), C2 (violet), SB1 (red), and SB2 (wine). Dotted lines represent AP half durations. (E) Scatter plots showing a statistical overview of the half duration in controls (C1 and C2) and patients (SB1 and SB2). \**p*<0.05 and \*\**p*<0.01 (nonparametric Mann–Whitney U test for pairwise comparisons). Color and symbol coding: C1, green diamond; C2, violet circle; SB1, red square; SB2, wine down-pointing triangle. Sample size: *n*=8 in C1, *n*=46 in C2, *n*=12 in SB1, and *n*=20 in SB2. Lines and whiskers represent mean values and the 5–95 percentile, respectively.

Altogether, comprehensive electrophysiological analyses indicate that SB iPSC-derived cells exhibited functional neuronal properties. Intriguingly, two intrinsic membrane parameters, AP frequencies and half durations, differed between SB and control neurons. In addition, the gain of the relationship between AP frequency and injected current stimulus was 2-fold greater in the SB cell groups than in the control groups.

## Discussion

The excessive involuntary trigeminal motor activity of SB can lead to a broad range of clinical complications in oro-dental regions. Our previous study found an association between the genetic variant of *HTR2A* and SB (Abe et al., 2012). The use of hiPSCs derived from patients with a specific neurological disease allows the preparation of brain cells that contain the actual genetic information of the patients themselves (Bellin et al., 2012; Park et al., 2008; Richard and Maragakis, 2015; Robinton and Daley, 2012; Saporta et al., 2011). This strategy is a notable feature given that such cells have been technologically and ethically difficult to obtain in the past (Imaizumi and Okano, 2014). In this study, SB and control hiPSCs were directly differentiated toward ventral hindbrain neurons, with slight modifications from the previously reported differentiation method, to elucidate neurological phenotypes associated with SB. We performed simultaneous differentiations of all four lines with triplicate wells and characterized control (C2) cell line utilizing RT-qPCR following a time-dependent course. Electrophysiological recordings were performed to assess the functional state of hiPSCs in long-term cultures, and to determine functional differences between SB and control neurons in the early stage of neurogenesis. We found that neurons obtained from C2 cell line mature in a time-dependent manner, similar to those of human brain neurons. In the early stage of neurogenesis, SB iPSC-derived neurons displayed alterations in AP frequency, half duration, and gain of the firing frequency-current intensity relationship.

### Electrophysiological maturation of hiPSC-derived neurons *in vitro*

Intrinsic membrane properties are an interpreting key perspective of neuronal maturation. Numerous neuronal intrinsic membrane parameters show critical alterations in neurogenesis (Kang et al., 2017; Kopach et al., 2018; Lam et al., 2017; Prè et al., 2014; Weick, 2016). The RMP of C2 iPSC-derived neurons shifted towards a more negative potential throughout neurogenesis, settling at approximately −63.9 mV, which is comparable to that of an adult human brain neuron (Testa-Silva et al., 2014). Similar to developing rodent neurons (Zhang, 2004), the RMP of generated neurons was depolarized (approximately −52.2 mV) at DIV31–51, steadily decreasing to approximately −63.9 mV in C2 neurons at DIV92–111. As a consequence of higher ion channel density and more complex cell morphology, R_m_ also decreased during neurogenesis (Kang et al., 2017; Kopach et al., 2018; Lam et al., 2017; Prè et al., 2014; Weick, 2016). Identical to the neurons from human brain tissue (Moore et al., 2009; Testa-Silva et al., 2014), R_m_ of long-term cultured neurons was higher, approximately 1692.2 MΩ at DIV31–51, dynamically decreasing to approximately 442.6 MΩ at DIV92–111. The τ_m_ also declined over time, matching to that of previously reported hiPSC-derived neurons (Prè et al., 2014). In line with previous observations, the C_m_ of generated neurons increased during neuronal maturation (Kang et al., 2017; Kopach et al., 2018; Kopach et al., 2020; Lam et al., 2017; Weick, 2016). Furthermore, during maturation, the AP threshold of differentiated neurons gradually hyperpolarized with the AP waveform displaying broader amplitudes with shorter half durations; as comparable to previously reported conventions (Lam et al., 2017; Testa-Silva et al., 2014; Zhang, 2004).

Taken together, quantitative comparisons of intrinsic membrane parameters clearly showed that hiPSC-derived neurons obtained from C2 line mature to a constitutive neurophysiological level in long-term cultures.

### Increased excitability of SB iPSC-derived neurons

This study documents the increased excitability of SB iPSC-derived neurons at the early stage of neurogenesis. Previous case-control studies of neurological disorders have shown increased neuronal excitability of the mutant in both animal models and hiPSC-derived neurons, including increased AP firing and a narrower AP half duration in neurons of transgenic Alzheimer’s disease (Ghatak et al., 2019) and Fragile X syndrome mouse models (Luque et al., 2017; Routh et al., 2017), as well as in hiPSCs of Alzheimer’s disease (Ghatak et al., 2019). In conjunction with increased spikes, elicited as a result of current injection, the spike rate gain was also found to be increased in Fragile X syndrome and autism spectrum disorder mice (Luque et al., 2017; Zhang et al., 2014). hiPSC-derived neurons in SB showed a higher number of APs for current steps of 50, 60, 70, and 80 pA, with slope of the firing frequency-current injection relationship increasing 2-fold. Moreover, this increase in APs was accompanied by shorter AP half-duration, leading to increased neuronal excitability in SB patients in the early stage of neurogenesis. Alterations in AP firing frequency and half duration indicate that the neuronal input-output function is modified in SB neurons.

Nevertheless, we acknowledge that a major limitation of this study is variability in culturing conditions between cell lines and the limited number of the subjects investigated, which were only compared at the early stage of neurodevelopment. Whether found neuronal excitability is indeed the phenotype-specific to SB iPSC-derived neurons remains to be determined. Further studies, of neural cells derived from a larger number of case and control subjects at later stages of neurodevelopment should be conducted to validate the results.

Another limitation of this study was that we did not collect information on the induced neuronal populations. While we have observed that a certain proportion of neurons expressed *HTR2A* by RT-qPCR, we did not evaluate the electrophysiological status corresponding to 5-HT2ARs. Besides, several neurological diseases might be associated with specific neuronal population, such as GABAergic neurons (Berretta et al., 2015; Kelley et al., 2008; Fan et al., 2017). Therefore, it is desirable to develop a tool that can label specific types of neurons. Such a strategy should also be developed in the future.

Within the limitation mentioned above of this study, our findings suggest that increased excitability of SB iPSC-derived neurons might be associated with the pathological mechanisms of SB. To the best of our knowledge, this study is the first to report the phenotypes of SB patient-derived neurons. Furthermore, this study provides a quantitative framework for considering the heterogeneity of various electrophysiological parameters for iPSC-based disease modeling.

## Materials and Methods

### Ethics statement

The following protocol was approved by the Showa University Ethics Committee for Genome Research (approval no. 179) and the Juntendo University School of Medicine Ethics Committee (approval no. 2016117). The study protocol was in accordance with the 1964 Declaration of Helsinki and its later amendments or comparable ethical standards. Formal informed consent was obtained from each individual.

### hiPSC derivation and neuronal differentiation

The hiPSCs of two SB patients with the risk allele (C) of *HTR2A* SNP rs6313 (SB1 and SB2) and two unaffected controls without the risk allele (C1 and C2) established in the previous study (Hoashi et al., 2017) were used. Neuronal differentiation was performed as previously described (Hoashi et al., 2017) but with minor modifications. hiPSC lines were dissociated to form embryoid bodies and cultured on mitomycin C-treated SNL murine fibroblast feeder cells in standard human embryonic stem cell culture (hESC) medium (DMEM/F12; Sigma Aldrich, St. Louis, MO, USA) containing 20% knockout serum replacement (Thermo Fisher Scientific, Waltham, MA, USA), 2 mM L-glutamine, 0.1 mM non-essential amino acids (Sigma Aldrich), 0.1 mM 2-mercaptoethanol (Sigma Aldrich), 0.5% penicillin/streptomycin, and 4 ng/mL fibroblast growth factor 2 (FGF-2; PeproTech, Rocky Hill, NJ, USA) in an atmosphere containing 3% CO_2_; the medium was changed every other day. hiPSCs were pretreated for 5 days with 3 μM SB431542 (Tocris, Bristol, UK), 3 μM dorsomorphin (Sigma Aldrich), and 3 μM CHIR99021 (ReproCELL, Kanagawa, Japan). Embryoid bodies were seeded at a density of 10 cells/μL in KBM Neural Stem Cell medium (Kohjin-Bio, Saitama, Japan) with specifically selected growth factors and inhibitors at 4% O2 and 5% CO_2_; growth factors and inhibitors included 1× B-27 supplement (Thermo Fisher Scientific), 0.5% penicillin/streptomycin, 20 ng/mL FGF-2, 2 μM SB431542, and 3 μM CHIR99021. The neurosphere culture was started on DIV0; neurospheres were passaged at a density of 50 cells/μL on DIV7 and DIV14. The neurosphere culture medium contained the following additives: 10 μM Y-27632 (Fujifilm, Tokyo, Japan) on DIV0–7, and 100 ng/mL Shh-C24II (R&D Systems, Minneapolis, MN, USA) and 1 μM purmorphamine (Millipore, Burlington, MA, USA) on DIV1–21. On DIV21, neurospheres were replated on dishes coated with a poly-L-lysine solution and fibronectin and cultured in 5% CO_2_ at a density of 40 × 10^4^ cells/well in a 48-well plate, 150 × 10^4^ cells/well in a 12-well plate, or 300 × 10^4^ cells/well in a 6-well plate. The medium was changed to Neurobasal Plus Medium (Thermo Fisher Scientific) supplemented with 1× B-27 Plus supplement (Thermo Fisher Scientific), 0.5% penicillin/streptomycin, 20 ng/mL brain-derived neurotrophic factor (BDNF; BioLegend, San Diego, CA, USA), 20 ng/mL glial cell-derived neurotrophic factor (GDNF; Alomone Labs, Jerusalem BioPark, Israel), 0.2 mM ascorbic acid (Sigma Aldrich), 0.5 mM dbcAMP (Nacalai Tesque, Kyoto, Japan), 1 ng/mL transforming growth factor-β (TGF-β; BioLegend), 10 μM DAPT (Sigma Aldrich), and 3 μM CHIR99021. Half of the medium volume was replaced with fresh medium (including all supplements except CHIR99021) every 3 or 4 days.

### Reverse transcription and quantitative polymerase chain reaction (RT-qPCR)

Total RNA was isolated with trizol reagent (15596026; Thermo Fisher Scientific). cDNA was prepared using SuperScript First-Standard (Thermo Fisher Scientific) or the TaqMan Fast Advanced Master Mix (Thermo Fisher Scientific). qRT-PCR analyses were performed on a QuantStudio 7 Flex (Thermo Fisher Scientific). Reactions were carried out in duplicate, and data were analyzed using the comparative (ddCt) method. Custom primers for the *GAPDH* gene (PN4331182; Thermo Fisher Scientific), *NKX2.2* gene (PN4331182; Thermo Fisher Scientific), *EN1* gene (PN4331182; Thermo Fisher Scientific), *HOXC4* gene (PN4331182; Thermo Fisher Scientific), *Nestin* gene (PN4331182; Thermo Fisher Scientific), *βIII-tubulin* gene (PN4331182; Thermo Fisher Scientific) and *HTR2A* gene (PN4331182; Thermo Fisher Scientific) were used in the TaqMan assay.

### Patch-clamp recordings in cultured cells

For whole-cell patch-clamp recordings, individual coverslips with hiPSC-derived neurons from two SB patients and two unaffected controls were transferred in a recording chamber attached to the stage of an Olympus BX51WI upright microscope (Tokyo, Japan) equipped with an infrared DIC imaging system. Recordings were performed in a normal bath solution at a speed of 1–2 mL/min, maintained at room temperature (23-27°C). The normal bath solution contained 130 mM NaCl, 26 mM NaHCO_3_, 10 mM glucose, 3 mM KCl, 2 mM CaCl_2_, 2 mM MgCl_2_, and 1.25 mM NaH_2_PO_4_. The solution was oxygenated with a mixture of 95% O_2_ and 5% CO_2,_ to establish a pH of 7.4. Neurons with a compact cell body, typically two or more extended processes, were selected. Patch pipettes were made from borosilicate glass capillaries (GD-1.5; Narishige, Tokyo, Japan) using a P-97 puller (Sutter, Atlanta, GA). The internal solution of the pipette contained 10 mM KCl, 130 mM K-gluconate, mM HEPES, 0.4 mM EGTA, 2 mM MgCl_2_, 0.3 mM Na_2_-GTP, and 2 mM Mg-ATP, with a pH of 7.25 and 285– 300 mOsm. Pipette resistance ranged from 2.5 to 5.0 MΩ when the electrodes were filled.

Electrophysiological signals were recorded using a MultiClamp 700B amplifier (Molecular Devices, Sunnyvale, CA, USA), filtered at 10 kHz, and sampled at 20 kHz using a Digidata 1440A digitizer (Molecular Devices). Access resistance was monitored continuously throughout the experiments. Data were analyzed on a personal computer using pCLAMP 10.7 software (Molecular Devices), Origin 2019 (OriginLab, Northampton, MA, USA), and Microsoft Excel.

### Electrophysiological recording protocol

Following whole-cell attainment, the passive membrane properties of hiPSC-derived neurons were recorded. In all recorded cells, RMP was assessed immediately after membrane breakthrough. Calculated liquid junctional potentials between the pipette and bath solutions were −12 mV, with all data corrected accordingly. Rm, τ_m (_the time for the neuron membrane potential to reach 63% of its resting value), and C_m_ were measured using the −10 mV hyperpolarizing current step. For current-clamp recordings, voltage responses were evoked from a holding potential of −60 mV using 300 ms-long steps ranging from −60 pA to +150 pA in 10 pA intervals delivered at 0.5 Hz. Rheobase (R_h_), and the threshold, were calculated from the first evoked AP in response to a depolarizing current. AP amplitudes were determined from the RMP to the peak of the AP, while the AP half duration was defined as the time required to reach an AP at the half level.

### Neuronal AP type classification

‘Type 0 neurons’ did not fire APs and were excluded from this study. ‘Type 1 neurons’ fired one AP above 0 mV, followed by a plateau. As the reversal potential of cations, the absolute limit of 0 mV was chosen, with mature APs generally reaching and overshooting this reversal potential. ‘Type 2 neurons’ fired more than one AP above 0 mV at a maximal frequency below 10 Hz, while ‘Type 3 neurons’ fired APs above 0 mV at a maximal frequency of 10 Hz or more. Our ranking framework for functional types of neurons adopted a spectrum that corresponding to the neuronal maturation stage; thus, Type 1 neurons were termed as least mature, while Type 3 neurons were considered to be more mature and functionally active.

### Statistical analysis

Data are expressed as the mean ± s.e.m. unless specified otherwise, with *n* referring to the number of cells examined. Shapiro–Wilk test was used to analyze whether experimental data sets follow a normal distribution. If the data were normally distributed and the variances were equal, we used one-way ANOVA followed by a Tukey’s post-hoc test for pairwise comparisons. If data point did not pass these criteria, we used nonparametric Kruskal–Wallis one-way analysis of variance on ranks, followed by Mann–Whitney U test for pairwise comparisons. Correlation between variables was assessed using the Pearson’s correlation coefficient (*r*). Categorical variables were assessed using Chi-squared test. A *p* value less than 0.05 was considered statistically significant. Statistical analyses were performed with IBM SPSS Statistics 27 (IBM, Armonk, NY, USA) software.

## Abbreviations

SB: sleep bruxism
SNP: single nucleotide polymorphism
hiPSC: human induced pluripotent stem cell
*HTR2A*: serotonin 2A receptor gene
5-HT2AR: serotonin 2A receptor
DMEM: Dulbecco’s modified Eagle’s medium
RA: retinoic acid
Shh: sonic hedgehog
PM: purmorphamine
DIV: days *in vitro*
RMP: resting membrane potential
R_m_: membrane resistance
τ_m_: time constant
C_m_: membrane capacitance
APs: action potentials
R_h_: AP rheobase
BDNF: brain-derived neurotrophic factor
GDNF: glial cell-derived neurotrophic factor
TGF-β: transforming growth factor-β
s.e.m.: standard error of the mean

## Acknowledgments

We thank Dr. Hideyuki Okano, Professor of Department of Physiology, Keio University School of Medicine, Tokyo, Japan, for providing hiPSCs.

## Data availability statement

The data supporting the results of this work are openly available in figshare at: https://doi.org/10.6084/m9.figshare.13642643.

## Author contributions

T.I., W.A., and K.B. study concept and experimental design. A.K.S., and S.N. electrophysiological recordings, analysis of data, and data interpretation. K.N. RT-qPCR experiments and hiPSC cultures. A.K.S., S.N., K.N., Y.A., T.I., W.A., and K.B. preparation of the initial draft of the manuscript. All other authors have contributed to data interpretation and critically reviewed the manuscript. All authors approved the final version of the manuscript and agreed to be accountable for all aspects of the work to ensure that questions related to the accuracy or integrity of any part of the work are appropriately investigated and resolved.

## References

Abe, Y., Suganuma, T., Ishii, M., Yamamoto, G., Gunji, T., Clark, G. T., Tachikawa, T., Kiuchi, Y., Igarashi, Y. and Baba, K. (2012). Association of genetic, psychological and behavioral factors with sleep bruxism in a Japanese population. J. Sleep Res. 21, 289–296.

Andoh-Noda, T., Akamatsu, W., Miyake, K., Matsumoto, T., Yamaguchi, R., Sanosaka, T., Okada, Y., Kobayashi, T., Ohyama, M., Nakashima, K., et al. (2015). Differentiation of multipotent neural stem cells derived from Rett syndrome patients is biased toward the astrocytic lineage. Mol. Brain 8, 31.

Arias, B., Gutiérrez, B., Pintor, L., Gastó, C. and Fañanás, L. (2001). Variability in the 5-HT(2A) receptor gene is associated with seasonal pattern in major depression. Mol. Psychiatry 6, 239–242.

Beddis, H., Pemberton, M. and Davies, S. (2018). Sleep bruxism: an overview for clinicians. Br. Dent. J. 225, 497–501.

Bellin, M., Marchetto, M. C., Gage, F. H. and Mummery, C. L. (2012). Induced pluripotent stem cells: the new patient? Nat. Rev. Mol. Cell Biol. 13, 713–726.

Berretta, S., Pantazopoulos, H., Markota, M., Brown, C. and Batzianouli, E. T. (2015). Losing the sugar coating: potential impact of perineuronal net abnormalities on interneurons in schizophrenia. Schizophr. Res. 167, 18–27.

Bostwick, J. M. and Jaffee, M. S. (1999). Buspirone as an antidote to SSRI-induced bruxism in 4 cases. J. Clin. Psychiatry 60, 857–860.

Carra, M. C., Macaluso, G. M., Rompré, P. H., Huynh, N., Parrino, L., Terzano, M. G. and Lavigne, G. J. (2010). Clonidine has a paradoxical effect on cyclic arousal and sleep bruxism during NREM sleep. Sleep 33, 1711–1716.

Ellison, J. M. and Stanziani, P. (1993). SSRI-associated nocturnal bruxism in four patients. J. Clin. Psychiatry 54. 432–434.

Fan, X., Qu, F., Wang, J. J., Du, X. and Liu, W. C. (2017). Decreased γ-aminobutyric acid levels in the brainstem in patients with possible sleep bruxism: A pilot study. J. Oral Rehabil. 12, 934–940.

Fujimori, K., Matsumoto, T., Kisa, F., Hattori, N., Okano, H. and Akamatsu, W. (2017). Escape from pluripotency via inhibition of TGF-β/BMP and activation of Wnt signaling accelerates differentiation and aging in hPSC progeny cells. Stem Cell Reports 9, 1675–1691.

Gerber, P. E. and Lynd, L. D. (1998). Selective serotonin-reuptake inhibitor-induced movement disorders. Ann Pharmacother. 32, 692–698.

Ghatak, S., Dolatabadi, N., Trudler, D., Zhang, X., Wu, Y., Mohata, M., Ambasudhan, R., Talantova, M. and Lipton, S. A. (2019). Mechanisms of hyperexcitability in Alzheimer’s disease hiPSC-derived neurons and cerebral organoids vs isogenic controls. Elife 8, e50333.

Guiard, B. P. and Di Giovanni, G. (2015). Central serotonin-2A (5-HT2A) receptor dysfunction in depression and epilepsy: the missing link? Front Pharmacol. 6, 46.

Higurashi, N., Uchida, T., Lossin, C., Misumi, Y., Okada, Y., Akamatsu, W., Imaizumi, Y., Zhang, B., Nabeshima, K., Mori, M. X., et al. (2013). A human Dravet syndrome model from patient induced pluripotent stem cells. Mol. Brain 6, 19.

Hoashi, Y., Okamoto, S., Abe, Y., Matsumoto, T., Tanaka, J., Yoshida, Y., Imaizumi, K., Mishima, K., Akamatsu, W., Okano, H., et al. (2017). Generation of neural cells using iPSCs from sleep bruxism patients with 5-HT2A polymorphism. J. Prosthodont. Res. 61, 242–250.

Holmes, C., Arranz, M. J., Powell, J. F., Collier, D. A. and Lovestone, S. (1998). 5-HT2A and 5-HT2C receptor polymorphisms and psychopathology in late onset Alzheimer’s disease. Hum. Mol. Genet. 7, 1507–1509.

Imaizumi, Y., Okada, Y., Akamatsu, W., Koike, M., Kuzumaki, N., Hayakawa, H., Nihira, T., Kobayashi, T., Ohyama, M., Sato, S., et al. (2012). Mitochondrial dysfunction associated with increased oxidative stress and α-synuclein accumulation in PARK2 iPSC-derived neurons and postmortem brain tissue. Mol. Brain 5, 35.

Imaizumi, Y. and Okano, H. (2014). Modeling human neurological disorders with induced pluripotent stem cells. J. Neurochem. 129, 388–399.

Inayama, Y., Yoneda, H., Sakai, T., Ishida, T., Nonomura, Y., Kono, Y., Takahata, R., Koh, J., Sakai, J., Takai, A., et al. (1996). Positive association between a DNA sequence variant in the serotonin 2A receptor gene and schizophrenia. Am. J. Med. Genet. 67, 103–105.

Kang, S., Chen, X., Gong, S., Yu, P., Yau, S., Su, Z., Zhou, L., Yu, J., Pan, G. and Shi, L. (2017). Characteristic analyses of a neural differentiation model from iPSC-derived neuron according to morphology, physiology, and global gene expression pattern. Sci. Rep. 7, 12233.

Kato, T., Thie, N. M., Huynh, N., Miyawaki, S. and Lavigne, G. J. (2003). Topical review: sleep bruxism and the role of peripheral sensory influences. J. Orofac. Pain 17, 191–213.

Kelley, J. M., Hughes, L. B. and Bridges, S. L. Jr. (2008). Does gamma-aminobutyric acid (GABA) influence the development of chronic inflammation in rheumatoid arthritis? J. Neuroinflammation 5, 1.

Kopach, O., Rybachuk, O., Krotov, V., Kyryk, V., Voitenko, N. and Pivneva, T. (2018). Maturation of neural stem cells and integration into hippocampal circuits − a functional study in an *in situ* model of cerebral ischemia. J. Cell Sci. 131, jcs210989.

Kopach, O., Esteras, N., Wray, S., Rusakov, D. A. and Abramov, A. Y. (2020). Maturation and phenotype of pathophysiological neuronal excitability of human cells in tau-related dementia. J. Cell Sci. 133, jcs241687.

Lam, R. S., Töpfer, F. M., Wood, P. G., Busskamp, V. and Bamberg, E. (2017). Functional maturation of human stem cell-derived neurons in long-term cultures. PLoS One 12, e0169506.

Lavigne, G. J., Khoury, S., Abe, S., Yamaguchi, T. and Raphael, K. (2008). Bruxism physiology and pathology: an overview for clinicians. J. Oral Rehabil. 35. 476–494.

Lee, Y. A. and Goto, Y. (2018). The Roles of Serotonin in Decision-making under Social Group Conditions. Sci. Rep. 8, 10704.

Lobbezoo, F., Ahlberg, J., Glaros, A. G., Kato, T., Koyano, K., Lavigne, G. J., de Leeuw, R., Manfredini, D., Svensson, P. and Winocur, E. (2013). Bruxism defined and graded: an international consensus. J. Oral Rehabil. 40, 2–4.

Luque, M. A., Beltran-Matas, P., Marin, M. C., Torres, B. and Herrero, L. (2017). Excitability is increased in hippocampal CA1 pyramidal cells of Fmr1 knockout mice. PLoS One 12, e0185067.

Moore, A. R., Filipovic, R., Mo, Z., Rasband, M. N., Zecevic, N. and Antic, S. D. (2009). Electrical excitability of early neurons in the human cerebral cortex during the second trimester of gestation. Cereb. Cortex 19, 1795–1805.

Park, I. H., Arora, N., Huo, H., Maherali, N., Ahfeldt, T., Shimamura, A., Lensch, M. W., Cowan, C., Hochedlinger, K. and Daley, G. Q. (2008). Disease-specific induced pluripotent stem cells. Cell 134, 877–886.

Prè, D., Nestor, M. W., Sproul, A. A., Jacob, S., Koppensteiner, P., Chinchalongporn, V., Zimmer, M., Yamamoto, A., Noggle, S. A. and Arancio, O. (2014). A time course analysis of the electrophysiological properties of neurons differentiated from human induced pluripotent stem cells (iPSCs). PLoS One 9, e103418.

Richard, J. P. and Maragakis, N. J. (2015). Induced pluripotent stem cells from ALS patients for disease modeling. Brain Res. 1607, 15–25.

Robinton, D. A. and Daley, G. Q. (2012). The promise of induced pluripotent stem cells in research and therapy. Nature 481, 295–305.

Routh, B. N., Rathour, R. K., Baumgardner, M. E., Kalmbach, B. E., Johnston, D. and Brager, D. H. (2017). Increased transient Na^+^ conductance and action potential output in layer 2/3 prefrontal cortex neurons of the fmr1^-/y^ mouse. J. Physiol. 595, 4431–4448.

Saporta, M. A., Grskovic, M. and Dimos, J. T. (2011). Induced pluripotent stem cells in the study of neurological diseases. Stem Cell Res. Ther. 2, 37.

Sateia, M. J. (2014). International classification of sleep disorders-third edition: highlights and modifications. Chest 146, 1387–1394.

Testa-Silva, G., Verhoog, M. B., Linaro, D., de Kock, C. P. J., Baayen, J. C., Meredith, R. M., de Zeeuw, C. I., Giugliano, M. and Mansvelder, H. D. (2014). High bandwidth synaptic communication and frequency tracking in human neocortex. PLoS Biol. 12, e1002007.

Weick, J. P. (2016). Functional properties of human stem cell-derived neurons in health and disease. Stem Cells Int. 2016, 4190438.

Williams, J., McGuffin, P., Nöthen, M. and Owen, M. J. (1997). Meta-analysis of association between the 5-HT2a receptor T102C polymorphism and schizophrenia. EMASS Collaborative Group. European Multicentre Association Study of Schizophrenia. Lancet 349, 1221.

Xu, H., Guan, J., Yi, H. and Yin, S. (2014). A systematic review and meta-analysis of the association between serotonergic gene polymorphisms and obstructive sleep apnea syndrome. PLoS One 9, e86460.

Zhang, Y., Bonnan, A., Bony, G., Ferezou, I., Pietropaolo, S., Ginger, M., Sans, N., Rossier, J., Oostra, B., LeMasson, G., et al. (2014). Dendritic channelopathies contribute to neocortical and sensory hyperexcitability in Fmr1(-/y) mice. Nat. Neurosci. 17, 1701–1709.

Zhang, Z. W. (2004). Maturation of layer V pyramidal neurons in the rat prefrontal cortex: intrinsic properties and synaptic function. J. Neurophysiol. 91, 1171–1182.

